# Structural insights into the recognition of histone H3Q5 serotonylation by WDR5

**DOI:** 10.1101/2020.12.24.424296

**Authors:** Jie Zhao, Wanbiao Chen, Yinfeng Zhang, Fan Yang, Nan Shen, Xuan Zhang, Xi Mo, Jianye Zang

## Abstract

Serotonylation of histone H3Q5 (H3Q5ser) is a recently identified posttranslational modification of histones that apparently acts as a permissive marker for gene activation in synergy with H3K4me3 during neuronal cell differentiation. However, any proteins which specifically recognize H3Q5ser remain unknown. Here, we discovered that WDR5 interacts with the N-terminal tail of histone H3 and functions as a ‘reader’ for H3Q5ser. Crystal structures of WDR5 in complex with H3Q5ser and H3K4me3Q5ser peptides revealed that the serotonyl group is accommodated in a shallow surface pocket of WDR5. Experiments in neuroblastoma cells demonstrate that WDR5 colocalizes with H3Q5ser in the promoter regions of cancer-promoting genes, where it promotes gene transcription to induce cell proliferation. Thus, beyond revealing a previously unknown mechanism through which WDR5 reads H3Q5ser to activate transcription, our study suggests that this WDR5-H3Q5ser mediated epigenetic regulation apparently promotes tumorigenesis.

## Introduction

Nucleosomes comprise 147 bp of DNA, wrapped around core histone octamers; these are the fundamental subunits of the chromatin that carries both genetic and epigenetic information to control developmentally and environmentally appropriate gene transcription and genomic regulation(*1*). Posttranslational modifications on histones (hPTM) are now understood to exert major regulatory impacts on diverse biological processes. A variety of hPTMs such as methylation, acetylation, phosphorylation, and ubiquitination, has been identified to be epigenetic marks driving gene regulatory networks controlling cell fate decisions. hPTMs are dynamically regulated by ‘writer’ and ‘eraser’ enzymes, and are recognized by dozens of ‘reader’ modules, and these proteins variously coordinate to specify many distinct biological outcomes(*2–4*). Owing to developments in mass spectrometry technologies, new histone modifications such as crotonylation, β-hydroxybutyrylation, lactylation, and serotonylation, have been reported to activate gene transcription, and these modifications have been functionally implicated in physiological responses to metabolic stress and neuronal cell differentiation, among many others(*5–8*).

Serotonin is a neurotransmitter that activates intracellular signaling pathways through cell surface receptors(*9, 10*). Additionally, serotonin can covalently modify and thereby regulate the functions of target proteins, including fibronectin, small GTPase, and Rac1, upon transamidation catalyzed by tissue transglutaminases (TGMs)(*11–13*). Although histone proteins are known to be suitable substrates for TGMs *in vitro*, it has been unclear for more than a decade whether histones are serotonylated by TGMs *in vivo*(*14*). A recent study showed that TGM2 can catalyze serotonylation of the Q5 position of histone H3; the resultant H3Q5ser are frequently present alongside trimethylation of histone H3K4 (H3K4me3). Of particular note, H3 histones doubly modified with K4me3/Q5ser marks were implication in the transcription activation of genes that function in neuronal cell differentiation and in the development of the central nervous system(*7*).

Although H3Q5ser has been proposed as permissive modifications thought to generally promote gene transcription, no ‘reader’ proteins which specifically recognize H3Q5ser to mediate gene activation have been confirmed experimentally. Nevertheless, 77 proteins were shown to associate with histone H3 in the presence of Q5ser and K4me3 dual modifications, including WD repeat-containing protein 5 (WDR5)(*7*). WDR5 functions as the core subunit of the mixed lineage leukemia (MLL) complex, which catalyzes the methylation of H3K4 upon interaction with the N-terminal tail of histone H3(*15*). Previous studies have implicated WDR5-mediated transcriptional activity in tumor growth, proliferation, differentiation(*16*), and metastasis(*17*). Also, there are reports that WDR5 binds the promoters of neuronal genes to regulate the transcription of genes affecting cell differentiation(*18*). Given its previously shown interaction with H3Q5ser, and considering the reported neuronal-development impacts of both the WDR5 protein and H3Q5ser, it is possible that WDR5 may be a ‘reader’ protein for H3Qser that deciphers epigenetic signals to activate neurodevelopment-related genetic programs.

In this study, we identified WDR5 as a ‘reader’ of histone H3Q5ser and demonstrated that serotonylation of Q5 significantly increases the binding affinity of WDR5 for histone H3. We elucidated the atomic mechanism through which WDR5 specifically recognizes H3Q5ser based on two crystal structures of WDR5 in complex with H3Q5ser or H3K4me3Q5ser peptide. Extending these structural insights, we showed that disrupting WDR5’s capacity to read the H3Q5ser restricts the proliferation of neuroblastoma cells. Thus, our study demonstrates how WDR5 reads H3Q5ser and shows how this recognition and subsequent occupancy at selected loci activates target gene transcription. Finally, the fact that multiple genes targeted for transcriptional activation based on this WDR5/H3Q5ser mechanism are known cancer-promoting genes may help explain the progression of neuroblastoma and could support the development of innovative anti-tumor therapeutics.

## RESULTS

### Identification of WDR5 as a reader of H3Q5ser

Serotonylation of histone H3Q5 (H3Q5ser, **Fig. 1a**) is catalyzed by TGM2 and is now known to mediate permissive gene expression during neuronal development. WDR5, a well-known H3 binding protein, was previously suggested as a possible reader protein of H3Q5ser because it was pulled-down in an assay using a peptide bearing an H3K4me3Q5ser modification as the bait(*7*). Therefore, we conducted streptavidin pull-down assays with purified recombinant human WDR5 (containing residues 22-334; named WDR5^22-334^) using synthetic biotinylated histone H3 peptides as interaction substrates; these peptides comprised the N-terminal 14 amino acids of H3 and carried either no modification or modifications including trimethylated K4 (K4me3) and/or serotonylated Q5 (Q5ser) (**Table S1**). We found that WDR5^22-334^ interacts with four types of H3 peptides (H3Q5ser, H3K4me3Q5ser, H3K4me3 as well as unmodified H3 peptides) (**Fig. 1b,** upper panel). Notably, serotonylation on Q5 strongly enhances the binding affinity of WDR5 to the H3 peptide, regardless of the presence or absence of K4me3 (**Fig. 1b,** upper panel).

**Fig. 1:**
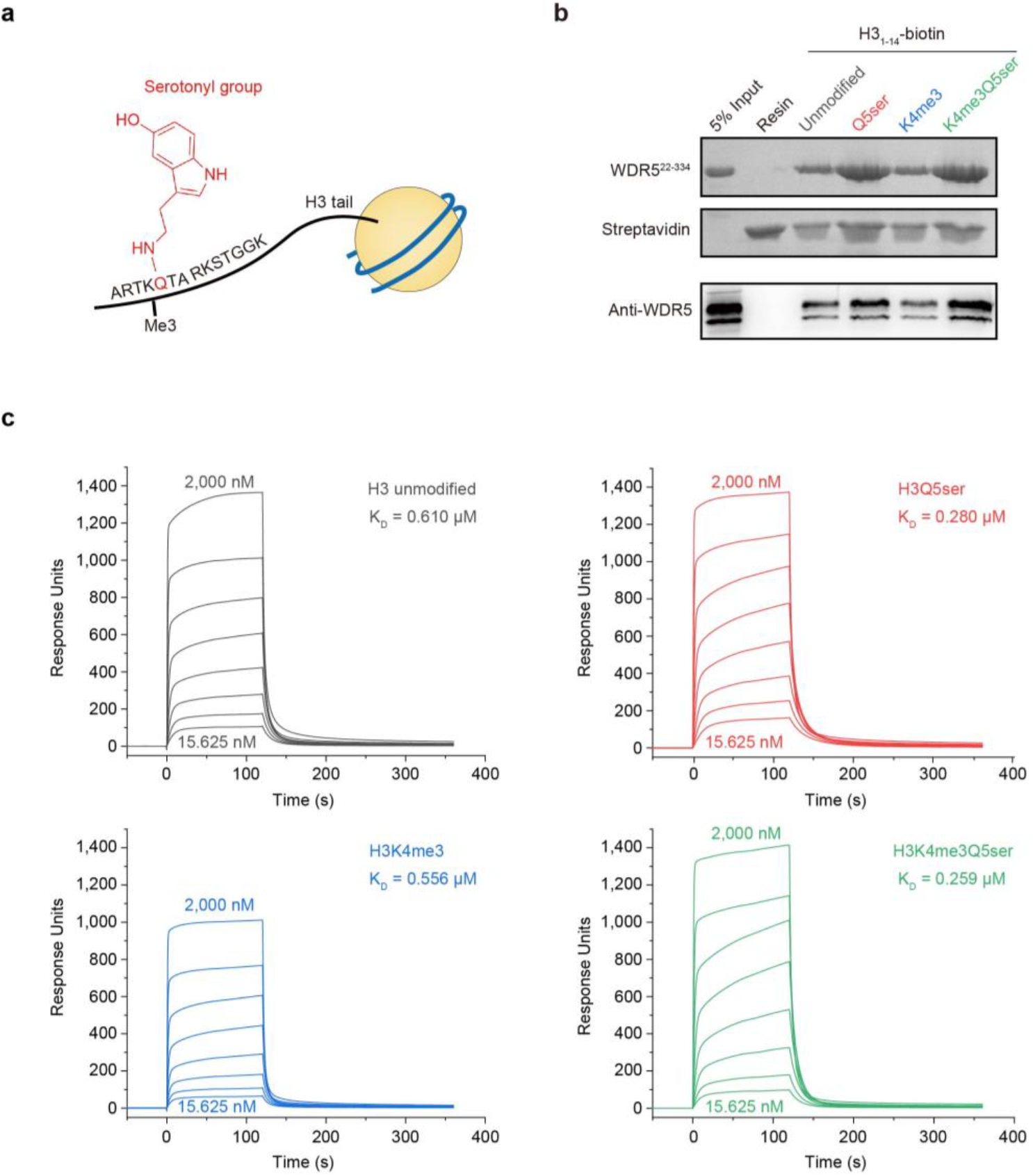
Serotonylation of H3Q5 facilitates WDR5 binding to H3. **a,** Chemical structure of serotonyl group covalently attached to the Q5 residue of histone H3. The histone octamer core and the wrapped DNA of the nucleosome are respectively shown as a solid yellow cycle and a blue line. The N-terminal 1-14 residues of H3 are indicated by their single-letter codes, with Q5 being colored red and the others colored black. Trimethylation on the K4 residue is indicated by “Me3”. The chemical structure of the serotonyl group on Q5 is shown in red. **b,** Streptavidin pull-down assay of recombinant WDR5^22-334^ (upper panel) or HEK 293T cell lysates overexpressing WDR5 (lower panel), with various biotinylated H3_1-14_ variants, with or without K4 trimethylation or Q5 serotonylation as indicated. **c,** SPR analysis to measure the binding affinity of WDR5^22-334^ with an unmodified H3 (gray) substrate peptide or H3 variant peptides bearing modifications including H3K4me3 (blue), H3Q5ser (red), or dual H3K4me3Q5ser (green).

To confirm the interaction of WDR5 with H3Q5ser, synthetic modified H3 peptides were incubated with cell lysates from HEK 293T cells that transiently over-expressed full-length human WDR5. Western blotting of the pulled-down proteins indicated that full-length WDR5 also binds to H3Q5ser/H3K4me3Q5ser bearing peptides with stronger affinity than for peptides bearing H3/H3K4me3 (**Fig. 1b,** lower panel). We also conducted surface plasmon resonance (SPR) assays to quantify the binding affinity of WDR5^22-334^ to the H3 peptides. WDR5^22-334^ binds to H3Q5ser peptide with the equilibrium dissociation constant (K_D_) of 0.280 μM, which is approximately 2.1 times lower than that of the unmodified H3 (K_D_ = 0.610 μM, **Fig. 1c** and **Table 2**). WDR5^22-334^ shows similar affinities with both H3Q5ser and H3K4me3Q5ser peptides, showing that the trimethylation of K4 has no obvious noticeable impacts on the interaction of serotonylated Q5 with WDR5. Taken together, these SPR and pull-down assays show that WDR5-H3 binding is strongly enhanced by the presence of a serotonylation modification to the H3 peptide’s Q5 residue.

**Table 2.** Binding affinity of wild type or mutant WDR5^22-334^ with various H3 peptides determined by SPR.

| H3<sub>1-14</sub> peptide | K<sub>D</sub> values (μM) |  |  |
| --- | --- | --- | --- |
|  | WDR5<sup>22-334</sup> |  |  |
|  | WT | N130A | Y131A |
| H3 | 0.610 | 0.664 | 0.761 |
| H3Q5<sub>ser</sub> | 0.280 | 0.418 | 0.567 |
| H3K4me3 | 0.556 | 0.774 | 1.087 |
| H3K4me3Q5<sub>ser</sub> | 0.259 | 0.517 | 0.947 |

### Overall structure of WDR5^22-334^ in complex with serotonylated H3 peptides

To further investigate the underlying molecular mechanism through which H3Q5ser interacts with WDR5, the crystal structures of WDR5^22-334^ in complex with H3Q5ser or H3K4me3Q5ser peptide were solved, both at 1.6 Å resolution (**Table 1**). In these two structures, the overall structures of WDR5^22-334^ are almost identical, with a root-mean-square distance (RMSD) value of 0.1 Å and both exhibiting the typical seven-bladed β-propeller structure of WD40 repeat proteins (**Fig. 2a**). Based on the electron density maps, the N-terminal residues 1-7 of these two peptides can be clearly traced, whereas residues 8-14 are disordered and not visible. Interpretable electron density was observed for the serotonyl group attached to Q5 in both structures, which was modeled unambiguously (**Fig. 2b, c**). Interestingly, no electron density was observed for the trimethyl group of K4 in the structure of the WDR5/H3K4me3Q5ser complex, likely because the side chain of K4me3 is flexible and protrudes towards the solvent.

**Table 1.**
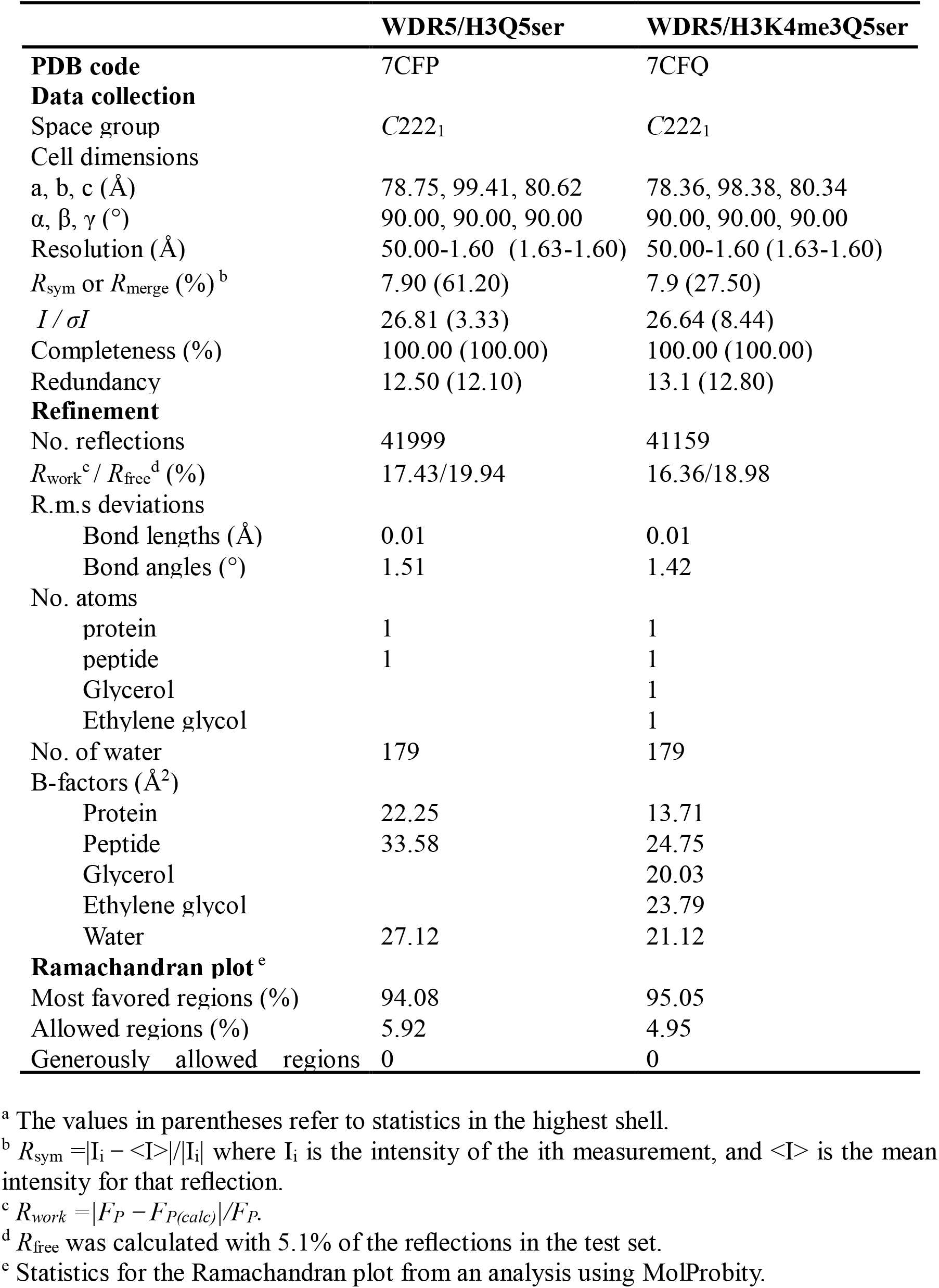
Data collection and refinement statistics.

**Fig. 2:**
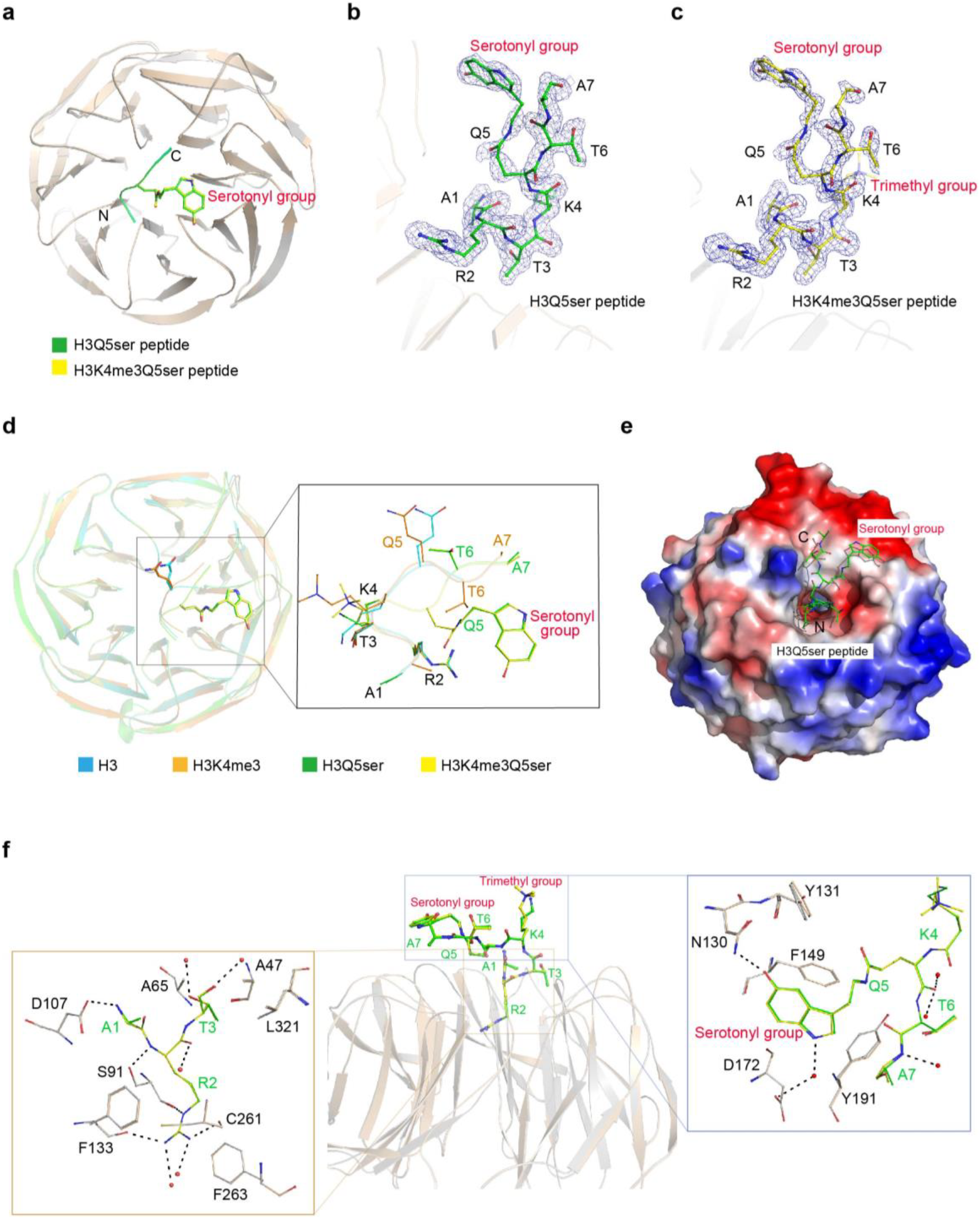
Crystal structures of WDR5^22-334^ in complex with either H3Q5ser or H3K4me3Q5ser peptide. **a,** Overall structures of WDR5^22-334^ in complex with either H3Q5ser or H3K4me3Q5ser peptide. The structures of WDR5^22-334^ are shown in a cartoon diagram; WDR5^22-334^/H3Q5ser is colored in wheat and WDR5^22-334^/H3K4me3Q5ser is colored in gray. The H3Q5ser (green) and H3K4me3Q5ser (yellow) peptides are shown in a cartoon diagram, with the serotonyl group on Q5 shown as sticks. **b and c**, Omit Fo – Fc electron density map of the H3Q5ser (**b**) and H3K4me3Q5ser (**c**) peptides, contoured at the 2.5 σ level. The peptides are shown as sticks and colored green and yellow (following color scheme from **a**). Trimethyl and serotonyl groups are indicated with red labels. **d,** Structural comparison of complexes including WDR5^22-334^/H3 (cyan, PDB code: 2H9M), WDR5^23-334^/H3K4me3 (Orange, PDB code: 2H6Q), WDR5^22-334^/H3Q5ser (Green), and WDR5^22-334^/H3K4me3Q5ser (Yellow). Enlarged view in the right panel shows binding-induced conformational changes in the peptides. **e,** Electrostatic potential surface view of WDR5 in complex with the H3Q5ser peptide. The peptide is shown as sticks. **f,** The interaction of H3Q5ser (green) and H3K4me3Q5ser (yellow) peptides with WDR5^22-334^. Amino acid residues of WDR5 involved in the peptide interaction are shown as sticks; these are colored wheat in WDR5^22-334^/H3Q5ser and colored gray in WDR5^22-334^/H3K4me3Q5ser. The enlarged views in the left and right panels show the detailed interactions between the H3 peptides and WDR5. Hydrogen bonds are indicated as dashed lines.

Alignment of the WDR5^22-334^/H3Q5ser and WDR5^22-334^/H3K4me3Q5ser complex structures with those of the WDR5/H3 (PDB code: 2H9M) complex and the WDR5/H3K4me3 complex (PDB code: 2H6Q) revealed that the overall structure of WDR5 in these structures is almost identical (**Fig. 2d**). The orientations of the first four resides 1-4 of histone H3 in these structures are also similar, whereas residues 5-7 undergo large conformational changes. Specifically, the side chain of Q5ser is flipped to the opposite side, thusly positioning the serotonyl group in a shallow hydrophobic surface pocket (**Fig. 2d, e**).

In both the WDR5^22-334^/H3Q5ser and WDR5^22-334^/H3K4me3Q5ser complex structures, the histone H3 N-termini are accommodated in the binding cleft of WDR5 via a series of hydrogen bonds and van der Waals contacts, findings similar to previously reported structures of WDR5 in complex with methylated histone H3K4 peptides (*19–22*). More specifically, R2 is anchored in a narrow central channel through cation-π interactions with F133 and F263. The side chains of both K4 and K4me3 are solvent exposed, and do not engage in any direct interactions with WDR5, helping to explain our findings from the SPR and streptavidin binding assays which indicated that the enhanced interaction of WDR5 with serotonylated histone H3 is independent of the methylation status of K4 (**Fig. 2e, f** and **Fig. 1c**).

### Specific recognition between WDR5 and serotonylated histone H3Q5

According to the complex structures, WDR5 specifically recognizes the serotonyl group through a network of hydrogen bonds and van der Waals contacts (**Fig. 3a, b**). In details, the amide group of the WDR5 N130 side chain forms a hydrogen bond with the hydroxyl group of serotonyl group attached to Q5. In addition, the WDR5 D172 side chain engages in water-mediated hydrogen bonding with the amide group of serotonin. The aromatic side chains of WDR5’s Y131, F149, and Y191 residues make van der Waals contacts with the hydrophobic moiety of the serotonyl group that help stabilize the WDR5-Q5ser interaction.

**Fig. 3:**
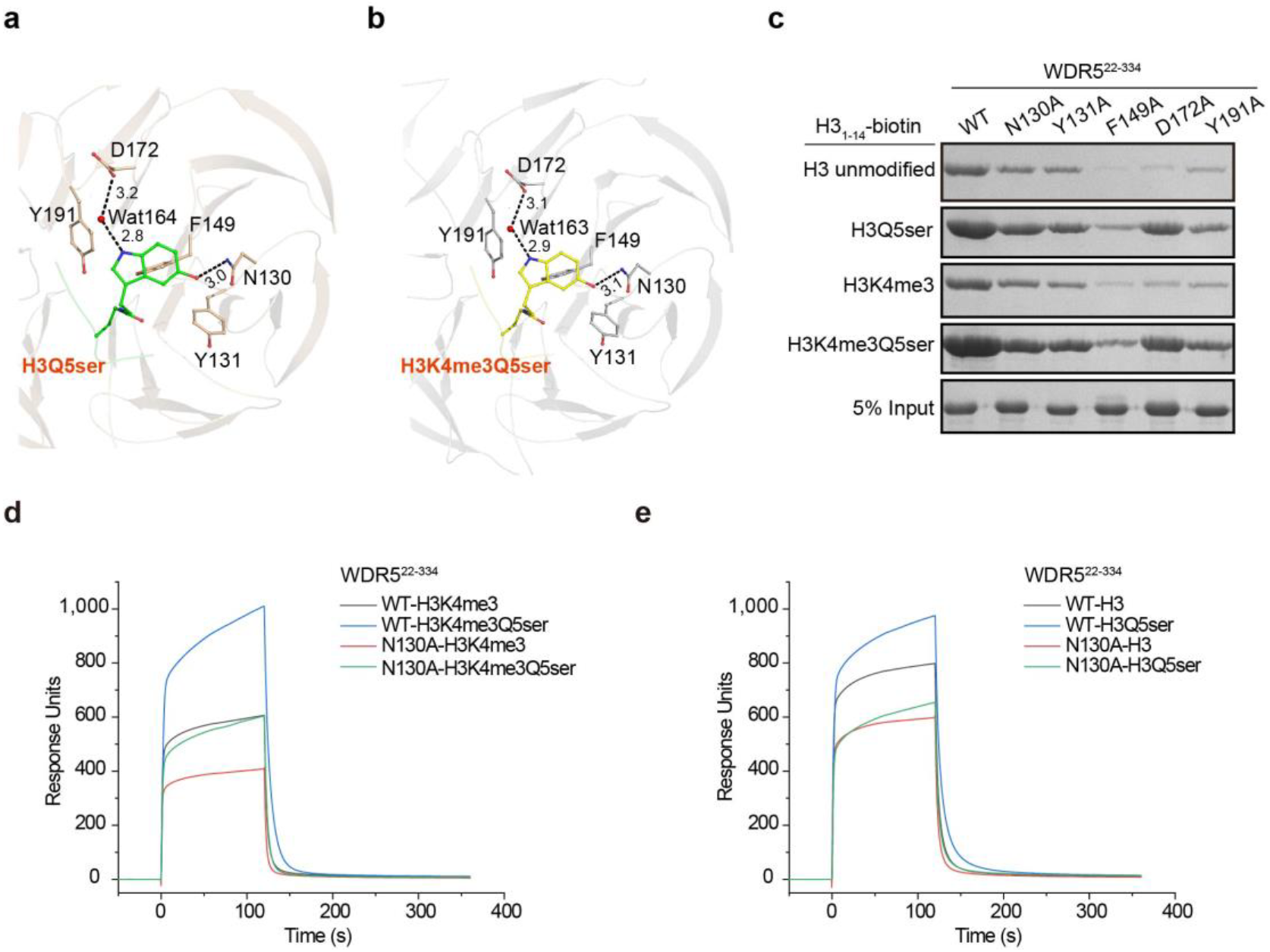
Recognition of serotonyl group by WDR5. **a and b**, Serotonyl group of H3Q5ser (a) and H3K4me3Q5ser (b) peptides bound to WDR5^22-334^. The hydrogen bonds are shown as dashed lines. Amino acid residues involved in serotonyl group binding are shown as sticks and are labeled in black. **c,** Streptavidin pull-down assays of wild type (WT) and five putative binding-deficient mutant variants of WDR5^22-334^ with the aforementioned biotinylated H3 peptides as substrates, as indicated. **d and e,** The binding of WDR5^22-334, WT^ and WDR5^22-334, N130A^ to H3 peptides, evaluated via SPR. Single-protein injections of WDR5^22-334, WT^ and WDR5^22-334, N130A^, all at 500 nM, over immobilized H3 or H3Q5ser peptides with (**d**) or without (**e**) K4 trimethylation.

To examine impacts of WDR5 residues involved in specific recognition of the serotonyl group, we generated a total of five WDR5 mutants to examine their interaction with histone H3 peptides. Circular dichroism spectroscopy and size-exclusion chromatography supported that all of the putative binding-deficient WDR5 mutant variants are soluble and properly folded (**Fig. S1**). Further, pull-down assays showed that the F149A, D172A, and Y191A WDR5 variants had obviously reduced interactions with all four types of H3 peptides compared to wild type WDR5 (*i.e.*, regardless of the presence or absence of Q5ser or K4me3). In contrast, the N130A and Y131A WDR5 variants had significantly reduced binding affinity for the serotonylated Q5-containing substrates but much less obvious difference for the unmodified H3 or H3K4me3 substrates. Collectively, these findings suggest that the functions of the N130 and Y131 residues are specific to the recognition of serotonylation modifications of histone H3Q5, whereas the F149, D172 and Y191 residues apparently functions in interactions with other histone H3 regions (**Fig. 3c**).

We next quantified the binding affinity of wild type WDR5 and WDR5 variants carrying the N130A or Y131A mutations for H3 peptide substrates by SPR experiments (**Fig. 3d, e, Table 2,** and **Fig. S2**). The equilibrium dissociation constants of mutant N130A (K_D_ = 0.664 μM) and wild type WDR5 (K_D_ = 0.610 μM) with unmodified H3 did not differ, whereas the K_D_ value of N130A (K_D_ = 0.418 μM) with H3Q5ser peptide was nearly 2 fold higher than with wild type WDR5 (K_D_ = 0.280 μM). Compared to wild type WDR5, the Y131A variant had a slightly increased K_D_ value (0.761 μM) for unmodified H3, but had a 2-fold higher K_D_ value (0.567 μM) for the H3Q5ser peptide substrate. These results confirm that both N130 and Y131 are responsible for the recognition of serotonylation of H3Q5.

Given reports that H3K4me3 frequently occurs alongside H3Q5ser modification, we next attempted to identify residues which may contribute to specific recognition of H3Q5ser modification rather than recognition of H3K4me3Q5ser dual modifications. The SPR data indicated that the binding affinity of the N130A variant (K_D_ = 0.517 μM) for the H3K4me3Q5ser peptide is significantly reduced compared to wild type WDR5 (K_D_ = 0.259 μM), whereas there was only slight difference in their respective binding with the H3K4me3 substrate, confirming the involvement of WDR5’s N130 residue in specific recognition of serotonylation at Q5 site. Similarly, the K_D_ value of the Y131A variant (0.947 μM) for H3K4me3Q5ser was increased by 3.7 folds compared to wild type WDR5 (K_D_ = 0.259 μM). However, the binding affinity of Y131A variant (K_D_ = 1.087 μM) for H3K4me3 peptide was also decreased by about 2 folds when compared to wild type WDR5 (K_D_ = 0.556 μM) (**Fig. 3d, e, Table 2,** and **Fig. S2**). These observations suggest that WDR5’s Y131 residue is also required for the recognition of Q5ser modification via hydrophobic interaction, but the presence of K4me3 modification interferes with the binding of the Y131A variant to H3K4me3 and H3K4me3Q5ser peptides in an unknown way. Taken together, these results identified N130 as a particularly impactful residue for the specific recognition of serotonyl group at H3Q5 site by WDR5, regardless of the presence or absence of K4me3.

### WDR5 regulates proliferation of neuroblastoma cells by recognition of H3Q5ser

To confirm the existence of H3Q5ser in neuroblastoma cells, lysates of the wild type neuroblastoma (SK-N-SH) cells were either directly loaded onto SDS-PAGE gel or first immunoprecipitated with an anti-histone H3 antibody, and were then immunoblotted with an anti-H3K4me3Q5ser antibody. H3Q5ser was detected in cell lysates; moreover it was detected in the anti-H3 antibody precipitates but not in control IgG precipitates (**Fig. 4a**), establishing that the Q5 site of H3 histones carry serotonylation modifications in neuroblastoma cells.

**Fig. 4:**
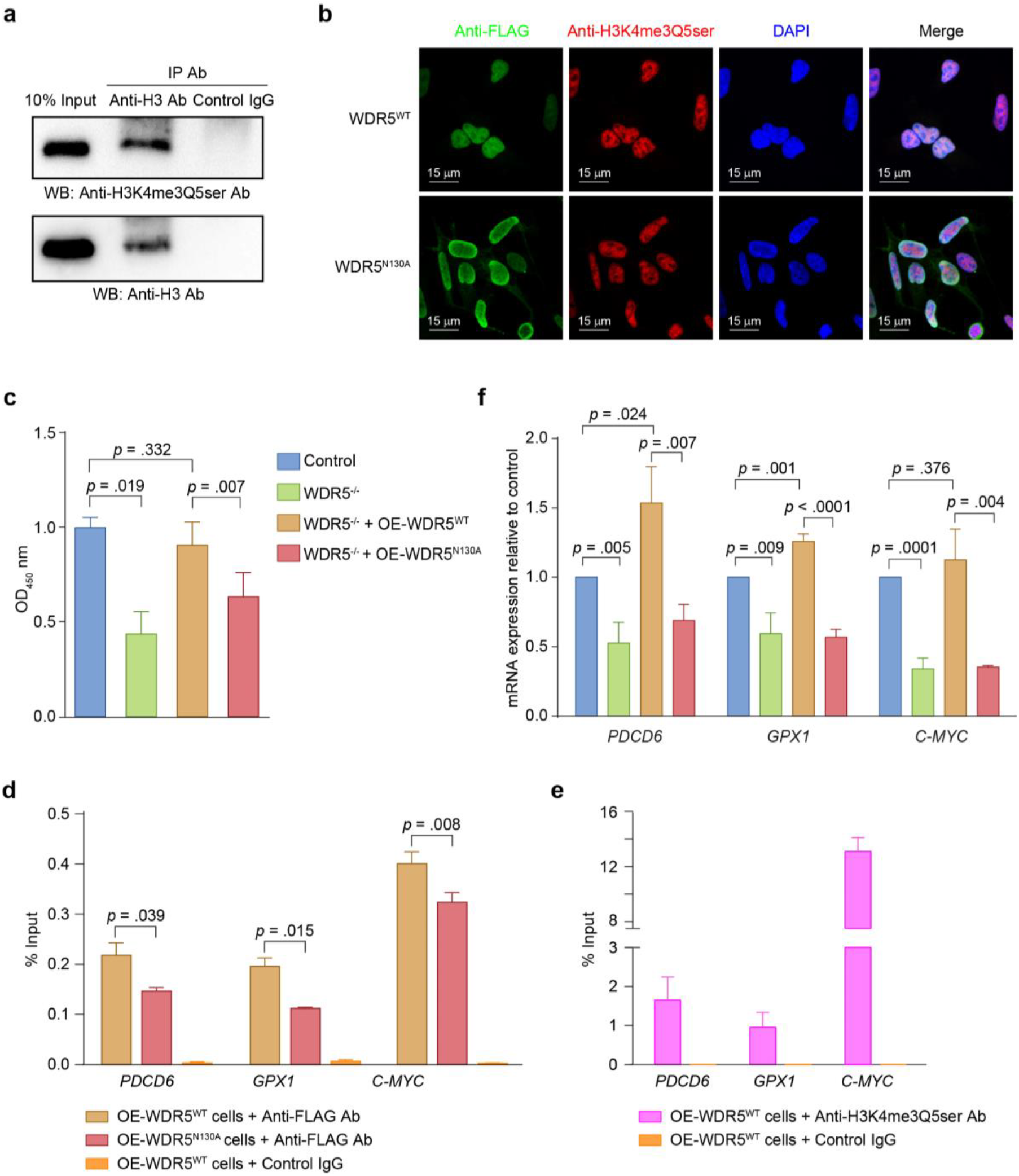
WDR5 regulates neuroblastoma cell proliferation by recognizing H3Q5ser. **a,** IP-WB analysis to detect the existence of H3Q5ser (upper panel) in human neuroblastoma SK-N-SH cells, both in directly loaded cell lysates and in anti-H3 antibody enriched precipitates. Histone H3 (lower panel) was visualized using an anti-H3 antibody (as a control). **b**, 3 × FLAG-tagged WDR5 (WT or with N130A mutation) were overexpressed in SK-N-SH cells. Co-localization of WT WDR5 or N130A mutant (green, anti-FLAG tag antibody) with H3Q5ser (red, anti-H3K4me3Q5ser antibody) in the SK-N-SH cell nuclei, as detected by immunofluorescence microscopy. **c,** Knockout of WDR5 via CRIPSR-Cas9 (WDR5^−/−^) significantly decreased the proliferation of SK-N-SH cells assessed by BrdU assays. The decreased cell proliferation phenotype was completely rescued upon complementation of the WDR5^−/−^ cells with WT but not N130A mutant WDR5. **d and e**, ChIP was performed with an anti-FLAG antibody (**d**, for WDR5) or an anti-H3K4me3Q5ser antibody (**e**, for H3Q5ser), in assays examining cells overexpressing 3 × FLAG tagged-WT WDR5 or N130A variant. qPCR analysis of the ChIP precipitates was performed to assess the co-occurrence of WDR5 occupancy and H3Q5ser modifications at the promoters of *PDCD6*, *GPX1*, and *C-MYC* genes shown to regulate tumor cell proliferation. **f**, mRNA expression levels of three cell proliferation-related genes in WDR5^−/−^ SK-N-SH cells or WDR5^−/−^ cells complemented with WT or N130A mutant WDR5, as assessed by qPCR. Data are presented as the mean ± standard deviation, calculated from three independent experiments, with the *p* value calculated by Student’s *t*-test as indicated. WT, wild type; OE, overexpression.

To further explore the association of WDR5 with H3Q5ser in cells, a 3 × FLAG tagged WDR5 fusion was expressed in SK-N-SH cells and both WDR5 and H3Q5ser were monitored by confocal microscopy after staining with anti-FLAG tag and anti-H3K4me3Q5ser antibodies. Both WDR5 (green) and H3Q5ser (red) were exclusively localized to neuroblastoma cell nuclei, and their signals were extensively colocalized (**Fig. 4b**). In contrast, much reduced co-localization was evident for cells expressing the N130A variant of WDR5, which recall has a significantly decreased capacity to interaction with H3Q5ser (**Fig. 3c, d**). Intriguingly, we observed that the N130A variant tended to localize at the nuclear membrane, and a very small portion of mutant proteins localized at the cytoplasm which was not observed for wild type WDR5 proteins; this finding suggests the possibility that H3Q5ser recognition by WDR5 may promote the translocation of WDR5 into the nucleus.

To explore potential pathological impacts of the WDR5-H3Q5ser interaction in neuroblastoma, we used CRIPSR/Cas9 to generate WDR5 knockout SK-N-SH cells. BrdU cell proliferation assays showed that WDR5 knockout (WDR5^−/−^) caused significantly decreased neuroblastoma cell proliferation compared to unedited cells (**Fig. 4c**). Note that these results are consistent with the previously reported clinical findings that high levels of WDR5 are correlated with poor prognosis of neuroblastoma patients(*23*). The decreased cell proliferation phenotype was completely rescued upon complementation of the WDR5^−/−^ cells with wild type WDR5. Moreover, we found that complementation with the N130A variant of WDR5 could only partially rescue the neuroblastoma cell proliferation. Thus, our results demonstrate that WDR5 functions to promote the proliferation of neuroblastoma cells and indicate that this proliferation-promoting impact of WDR5 is exacerbated by the presence of serotonylation modifications at the Q5 site of H3 histones.

WDR5 has been reported to bind to various promoter regions known to function in tumor maintenance(*24*), including genes that regulate tumor cell proliferation such as *PDCD6*, *GPX1*, and *C-MYC*(*25–27*). We therefore determined whether there is enrichment for WDR5 occupancy and/or H3Q5ser modifications in the regulatory regions of these loci in neuroblastoma cells. ChIP was performed with an anti-FLAG antibody (for WDR5) or an anti-H3K4me3Q5ser antibody (for H3Q5ser), in assays examining cells overexpressing wild type 3 × FLAG tagged-wild type WDR5 or N130A variant. qPCR analysis of the ChIP precipitates revealed the co-occurrence of WDR5 occupancy and H3Q5ser modifications at the promoters of *PDCD6*, *GPX1*, and *C-MYC* (**Fig. 4d, e)**. Importantly, we observed significantly reduced occupancy of the N130A variant of WDR5 at the promoters of these three loci compared to wild type WDR5, results establishing that H3Q5ser recognition promotes the recruitment of WDR5 to promoter of cancer-promoting genes.

To further confirm that WDR5 regulates gene expression by recognizing H3Q5ser, mRNA levels of the *PDCD6*, *GPX1*, and *C-MYC* genes were measured via qPCR. As expected, the expression of all three genes was significantly reduced in WDR5^−/−^ neuroblastoma cells, but the expression levels could be rescued to the levels of unedited cells upon complementation with wild type WDR5; complementation with the N130A WDR5 variant could only slightly rescue these expression levels, if not at all (**Fig. 4f**). Collectively, these experiments in neuroblastoma cells demonstrate that WDR5 binds to H3Q5ser present in gene promoter regions and activates gene transcription to facilitate cell proliferation.

## Discussion

Serotonylation of H3Q5 is the first endogenous monoaminyl modification of histones that has been identified(*7*). However, the reader proteins for H3Q5ser and its biological functions remain largely unknown. In the present study, we discovered that WDR5 functions as a H3Q5ser reader. We show that WDR5 preferentially binds serotonylated H3 over unmodified or H3K4me3 peptides, and both biochemical and structural studies—including solving two crystal structures for WDR5^22-334^/H3Q5ser and WDR5^22-334^/H3K4me3Q5ser complexes—revealed the residues and atomic interactions through which WDR5 recognizes H3Q5ser. Specifically, WDR5’s N130 residue mediates binding with the serotonyl group by contributing a hydrogen bond interaction.

WDR5 is considered to be an oncogene because its overexpression is related to poor prognosis in various human cancers such as neuroblastoma, colon cancer, prostate cancer, gastric cancer, breast cancer, lung cancer, and bladder cancer(*23, 28–33*). The underlying oncogenic mechanisms of WDR5 have to date been largely attributed to its interaction with MLL1 and its subsequent catalysis of trimethylation at H3K4 sites(*15*). In addition to MLL1, WDR5 can be recruited to N-myc target gene promotors through its physical interaction with N-myc, doing so in a manner that can promote H3K4 trimethylation in neuroblastoma; this WDR5 recruitment results in elevated transcription of N-myc target genes to promote tumor formation(*23*).

A clinically relevant aspect of our study is our demonstration from experiments in neuroblastoma cells that WDR5 undergoes a specific interaction with histone H3Q5ser to promote its localization at the promoter regions of *PDCD6*, *GPX1*, and *C-MYC*, genes known to promote the proliferation of tumor cells including neuroblastoma (*25–27*). We show that disrupting the interaction between WDR5 and H3Q5ser trapped WDR5 protein at the nuclear membrane, preventing its nucleus accumulation, which may lead to the decrease of the occupancy of WDR5 at target gene promoters. Such decrease reduced the expression levels of target genes, and decreased the extent of neuroblastoma cell proliferation. Together these findings suggest that specific recognition of H3Q5ser by WDR5 should be understood as an activating mechanism for the proliferation of neuroblastoma cells.

Histone H3Q5ser frequent co-exists alongside H3K4me3 on euchromatin, and these dual modifications are known to support efficient recruitment of the TFIID complex and to be associated with “permissive gene expression”(*7*). Our work here with neuroblastoma cells also shows that recognition of H3Q5ser facilitates the recruitment of WDR5 to the chromatin (**Fig. 4b, d, and e**). These findings support that serotonylation of histone H3Q5 should be understood as an epigenetic mark that is correlated with transcriptional activation. Besides WDR5, there are several reader proteins for histone H3K4me3 (*e.g.*, TAF3, BPTF, and Spindlin-1) that have been found to interact with histone H3K4me3Q5ser peptides(*7*). Previous studies have shown that WDR5, TAF3, BPTF, and Spindlin-1 are involved in activating gene transcription through their interactions with histone H3K4me3(*15, 34–37*). It should be highly informative to determine whether these proteins can recognize H3Q5ser, whether they impact gene transcription and to dissect whether their regulatory functions are specifically impacted by the simultaneous presence of H3Q5ser and H3K4me3 marks at target loci. Moreover, our study reveals insights about the progression of neuroblastoma, and suggests the opportunity to target WDR5 and/or H3Q5ser-related regulatory networks for developing innovative antitumor therapies.

## Materials and Methods

### Reagents and cell cultures

The human neuroblastoma cell line SK-N-SH and the human embryonic kidney cell line (HEK 293T) were obtained from the Cell Bank of Chinese Academy of Sciences. The SK-N-SH and HEK 293T cells were cultured in 1:1 Eagle’s Minimum Essential Medium (EMEM) (ATCC) and Ham’s F12 medium (Thermo) supplemented with 15% fetal bovine serum (FBS) and Dulbecco’s Modified Eagle’s Medium (DMEM) supplemented with 10% FBS, and incubated at 37°C in a humidified incubator with 5% CO_2_.

The antibodies used in current study included anti-WDR5 (Abcam, ab178410); anti-Histone H3K4me3Q5ser (Millipore, ABE2580); anti-FLAG tag (Affinity, T0003); anti-Histone H3 (Immunoway, YM3038); anti-β-Actin (Sigma, A3854); anti-β-Tublin (Yeasen, 30303ES50); HRP-conjugated anti-rabbit IgG (CST, 7074S); and HRP-conjugated anti-mouse IgG (CST, 7076S).

### Protein expression and purification

The human WDR5 fragment 22-334 (WDR5^22-334^) was cloned into the pET-28a vector (that contains a TEV cleavable N-terminal 8 × His tag) and was overexpressed in *Escherichia coli* Rosetta2 strain in LB medium. Cells were induced with 0.4 mM IPTG and grown at 16 °C for an additional 20 h for protein expression. Cells were harvested and resuspended in lysis buffer containing 50 mM Tris-HCl, pH 7.5, 500 mM NaCl, 5% (v/v) glycerol, 5 mM imidazole, 1 mM phenylmethylsulfonyl fluoride (PMSF), 2 mM β-mercaptoethanol (β-ME), and 20 μg/mL DNase1 (shyuanye, 9003-98-9), and lysed with a high-pressure homogenizer. The cell lysate was centrifuged and the supernatant was incubated with Ni-NTA affinity resin (GE Healthcare). The target protein was eluted with lysis buffer supplemented with 300 mM imidazole. The 8 × His tag was removed by TEV protease and tag-free WDR5^22-334^ was further purified by gel filtration using a Superdex 75 (10/300) increase column (GE Healthcare) in 20 mM Tris-HCl, pH 7.4, 150 mM NaCl, 1 mM EDTA, and 0.5 mM TCEP. The eluted WDR5^22-334^ protein was collected and concentrated for further use. All WDR5^22-334^ mutants were generated by PCR-based site-directed mutagenesis and were purified using the same procedure as the WDR5^22-334^ protein.

### Streptavidin pull-down assay

Streptavidin pull-down assays were performed to verify the binding activity of recombinant WDR5^22-334^ or full-length WDR5 expressed in mammalian cells with serotonylated H3 peptides.

For recombinant WDR5^22-334^, 10 μM synthetic biotinylated H3 peptides were incubated with 1.15 μM purified tag-free WDR5^22-334^ proteins, WT or mutant, in 1 mL binding buffer (gel filtration buffer containing 0.2% NP-40) at 4 °C for 40 min. The reaction components were added to 15 μL streptavidin-agarose resin and incubated at 4 °C for another 40 min. The beads were washed four times with binding buffer, and boiled at 100 °C. The bound proteins were analyzed by SDS-PAGE followed by Coomassie brilliant blue staining.

For full-length WDR5 expressed in mammalian cells, HEK 293T cells were transiently transfected with pLVX-IRES-Puro-WDR5 plasmid by PEI (Polysciences, 23966-2) according to the manufacturer’s instructions. Forty hours after transfection, the cells were lysed with 25 mM Hepes, pH 7.5, 100 mM NaCl, 50 mM KCl, 2% Glycerol, 1 mM EDTA, 2 mM PMSF, complete protease inhibitor cocktail (Roche 04693124001), and 2% NP-40.

Cell lysates were incubated with 2 μM various biotinylated H3 peptides following the similar procedures described above. The bound proteins were detected with rabbit anti-WDR5 antibody (Abcam ab178410)

### Surface plasma resonance (SPR)

SPR experiments were performed on a Biacore T200 instrument (GE Healthcare). The four biotinylated H3 peptides were immobilized on channel FC2 and FC4 of two Sensor SA Chips, while channel FC1 and FC3 were used as blank controls. The free sites of the surface were blocked with 1 mM biotin. All measurements were carried out at 298 K in buffer containing 10 mM Hepes, pH 7.4, 150 mM NaCl, 3 mM EDTA, 5 mM β-ME, and 0.01% P-20. Recombinant WDR5^22-334^ proteins were diluted as eight different concentrations (2 μM, 1 μM, 500 nM, 250 nM, 125 nM, 62.5 nM, 31.25 nM, and 15.625 nM) and were injected into each channel at a flow rate of 30 μL/min. Each injection cycle comprised 120 s sample injection, 240 s buffer flow for dissociation, and 60 s 50 mM NaOH flow for regeneration. The experimental data were analyzed by Biacore T200 BiaEvaluation software using the steady state affinity binding model.

### Circular dichroism (CD)

Far-UV CD spectrum signals were detected with an Applied Photophysics Chirascan spectrometer. All measurements were carried out at 298 K in buffer containing 10 mM Tris-HCl, pH 7.4, and 50 mM NaCl, at wavelengths ranging from 190 nm to 260 nm. All samples were recorded for three times and each final spectra curve was the average of three scans.

### Crystallization, X-ray data collection, and structure determination

Purified WDR5^22-334^ was mixed with synthetic H3Q5ser or H3K4me3Q5ser peptide at a molar ratio of 1:5 for 2 h on ice before crystallization. The crystals of the WDR5^22-334^/H3Q5ser complex were grown in 0.1 M sodium citrate tribasic dihydrate pH 5.5; 22% PEG 3,350; and 0.1% n-Octyl-β-D-glucoside at 295 K. Crystals of the WDR5^22-334^/H3K4me3Q5ser complex were obtained in 0.15 M ammonium sulfate; 0.1 M MES pH 5.5; 25% PEG 4,000; and 2% PEG 3,350.

The X-ray diffraction datasets were collected at the National Facility for Protein Science in Shanghai (NFPS) at beamline BL18U1. All data were processed, integrated, and scaled using HKL2000(*38*).

The crystal structure of WDR5 (PDB code:2GNQ) was used as the searching model for molecular replacement by phaser(*39*) in Phenix(*40*). Structure refinement, manual model building, and structural analysis were carried out using REFMAC5(*41*) and Wincoot(*42*). Data collection and structure refinement statistics are summarized in Table 1. Figures were generated using Pymol (http://www.pymol.org).

### Lentivirus-mediated WDR5 knockout and overexpression

To generate WDR5 knockout SK-N-SH cells by CRISPR/Cas9-mediated gene editing, two independent guide sequences targeting WDR5 gene were designed (E-CRISP, http://www.e-crisp.org/E-CRISP/) and cloned into the LentiCRISPR-v2-BSD plasmid, which was derived from the lentiCRISPR v2 plasmid (Addgene Plasmid #52961) by replacing the puromycin gene with the blasticidin gene. The sgRNA sequences targeting WDR5 gene are as follows: sgRNA1: CTGGGACTACAGCAAGGGGA and sgRNA2: CAGAAACTACAAGGCCACAC. The guide RNA-containing lentiviral plasmid was then co-transfected with two packing plasmids psPAX2 and pMD2.G into HEK 293T cells to generate lentivirus. SK-N-SH cells were transduced with the lentivirus for 48 h, and were subjected to blasticidin selection at 3 μg/mL for 14 days to obtain stable WDR5 knockout (WDR5^−/−^) cell lines.

To obtain WDR5 overexpression cells, full-length cDNA encoding WDR5 was amplified from the human brain cDNA library and cloned into a pLL3.7-puro tagged cloning vector encoding a C-terminal 3 × FLAG tag. Mutagenesis of WDR5 N130A was carried out using a Q5 site-Directed MUtagenesis Kit (NEB, E0552S). Lentivirus packaging and infection were carried out as described above, either in control SK-N-SH cells or in WDR5^−/−^ cells.

### Cell proliferation assay

Cell proliferation was assayed using a Bromodeoxyuridine (BrdU) Cell Proliferation Assay Kit (CST, #6813) following the manufacturer’s instructions. Briefly, cells were seeded at a density of 4 × 10^3^ cells/well in a 96-well plate. 10 μM BrdU labeling solution was substituted and incubated for 4 h; cells were then fixed for 1 h and incubated with anti-BrdU for 1 h. After removal of the unbound BrdU, the cells were incubated with an HRP labeled secondary antibody for 30 min. TMB substrate solution was then added, and the absorbance at 450 nm was measured using a BioTek Synergy/2 microplate reader (BioTek)

### Immunoprecipitation (IP) and Western blotting

6 ×10^6^ SK-N-SH cells were lysed with RIPA lysis buffer (50 mM Tris-HCl, pH 7.4, 150 mM NaCl, 1 mM EDTA, 1% Triton X-100, 1% Sodium deoxycholate, and 0.1% SDS) supplemented with 1% protease and phosphatase inhibitor cocktail (Thermo) at 4 °C for 30 min. The cell lysates were immunoprecipitated with anti-H3 antibody or control IgG at 2 μg/mL and Protein G agarose beads at 4 °C for 1 h. Proteins bound to the agarose beads were separated on a 15% SDS-PAGE gel, transferred to a PVDF membrane, immunoblotted with an anti-H3K4me3Q5ser or anti-H3 antibody, and finally visualized using an enhanced chemiluminescence (ECL) substrate kit (Millipore) with an ImageQuant LAS 4000 mini densitometer (GE Healthcare Life Science).

### Immunofluorescence staining and confocal microscopy imaging

SK-N-SH cells were seeded onto 12 mm slides (Fisher) in a 24-well plate one day before immunofluorescence staining. Cells stably expressing 3 × FLAG-tagged full-length or N130A mutant WDR5 were fixed with 4% PFA in PBS at room temperature for 10 min, and then permeabilized with 0.5% Triton X-100 in PBS for 10 min. After blockage with 5% BSA and 5% goat serum in PBS at room temperature for 30 min, the cells were incubated with anti-Histone H3K4me3Q5ser (1:200 diluted in PBST) or anti-FLAG tag (1:1000 diluted in PBST) antibody for 1 h, followed by incubation with a fluorescently labeled secondary antibody (1:500 diluted in PBST) for 30 min. After DAPI staining, cells were dehydrated with ethanol and mounted with ProLong Diamond Antifade Mountant (Invitrogen). Images were acquired using a Leica SP8 confocal microscope.

### Quantitative PCR (qPCR)

Total RNA of various cell samples was extracted using TRIzol (Thermo). Hieff qPCR SYBR Green Master Mix (Yeasen) was used for mRNA quantification following the manufacturer’s instructions. Briefly, RNA reverse transcription was performed with 5 × PrimerScript RT Master Mix (Takara). qPCR was performed using a CFX Connect™ Real-Time System (Bio-Rad) under the following cycling condition: 95 °C for 2 min, 40 amplification cycles of 95 °C for 5 s and 60 °C for 30 s, followed by a final cycle of 95 °C for 5 s and 65 °C for 5 s. Relative expression levels of target genes were calculated using the 2^−ΔΔCT^ method, and the levels of these genes in control SK-N-SH cells were set at 1 for normalization. qPCR for each gene was performed in technical triplicates in three independent experiments. Sequences of the primer used are listed in Table S2.

### Chromatin immunoprecipitation and quantitative PCR (ChIP-qPCR)

ChIP assays were performed following the protocol of the Myers Lab at Stanford University (http://www.hudsonalpha.org/myers-lab/protocols/). Briefly, 2 × 10^7^ cells in culture dishes were fixed with 1% formaldehyde at room temperature for 10 min, and the fixation was stopped via addition of 0.125 M glycine at room temperature for 5 min. The cells were washed with PBS and scraped into cold lysis buffer (10 mM Tris-HCl, pH 7.4, 10 mM NaCl, 3 mM MgCl_2_, 0.5% NP-40). Chromatin pellets obtained by centrifugation were re-suspended with 0.8 ml of RIPA buffer (50 mM Tris-HCl, pH 7.5, 300 mM NaCl, 1.0% Triton X-100, 0.5% sodium deoxycholate, 0.1% SDS,) and sonicated at 25% amplitude for a total of 8 min (with intervals). Sonicated chromatin was then incubated with 2 μg of anti-Histone H3K4me3Q5ser, anti-FLAG tag antibody, or control IgG pre-coated on Dynabeads protein G (Life Technologies) at 4 °C overnight. Immunoprecipitants were washed twice with RIPA buffer, and washed an additional three times with LiCl washing buffer (100 mM Tris-HCl, pH 7.5, 500 mM LiCl, 1% NP-40, 1% sodium deoxycholate). Chromatin was then eluted by protease K digestion, was crosslink reversed at 65 °C overnight, and was purified using a QIAquick PCR purification kit (Qiagen). The ChIPed DNA was assessed qPCR as described above; primer information is listed in Table S3.

### Statistical analysis

The data from at least three independent experiments were expressed as the mean ± SD (standard deviation). Two-tailed Student’s *t*-tests were used to assess the paired samples. All graphs were generated using GraphPad PRISM 6.0 (GraphPad Software, Inc., USA). A *p* value < 0.05 was considered statistically significant.

## Acknowledgements

The authors would like to thank the staff at beamline BL18U1 of the Shanghai Synchrotron Radiation Facility of the National Facility for Protein Science in Shanghai for the assistance with data collection. This work was supported by the Strategic Priority Research Program of the Chinese Academy of Sciences (XDB37010105), the National Key Research and Development Program of China (2017YFA0503600, 2016YFA0400903), the Foundation for Innovative Research Groups of the National Natural Science Foundation of China (31621002), the National Natural Science Foundation of China (91853133 to JZ, 31700671 to XZ, and 82002662 to YZ), the USTC Research Funds of the Double First-Class Initiative (YD2070002015), the Fundamental Research Funds for the Central Universities (WK2070000150, WK2070000171) and Science and Technology Commission of Shanghai Municipality (18ZR1424500 to XM).

## Author contributions

Jianye Zang, Xi Mo, Xuan Zhang and Nan Shen provided the scientific direction and the overall experimental design for the studies. Jie Zhao performed the crystallization and biochemical experiments. Jie Zhao and Wanbiao Chen collected the X-ray diffraction data sets, and Xuan Zhang was responsible for the crystal structure determination. Wanbiao Chen, Yinfeng Zhang and Fan Yang constructed the cell lines and carried out the cell biological experiments. Jie Zhao, Wanbiao Chen, Nan Shen, Xuan Zhang, Xi Mo and Jianye Zang wrote the manuscript.

## Reference

1. K. Luger, Mader, A.W., Richmond, R.K., Sargent, D.F. and Richmond, T.J., Crystal structure of the nucleosome core particle at 2.8 angstrom resolution. Nature 389, 251–260 (1997).

2. T. Kouzarides, Chromatin modifications and their function. Cell 128, 693–705 (2007).

3. I.O. Torres, D.G. Fujimori, Functional coupling between writers, erasers and readers of histone and DNA methylation. Curr Opin Struc Biol 35, 68–75 (2015).

4. D.J. Patel, A Structural Perspective on Readout of Epigenetic Histone and DNA Methylation Marks. Cold Spring Harb Perspect Biol 8, (2016).

5. Z. Xie, D. Zhang, D. Chung, Z. Tang, H. Huang, L. Dai, S. Qi, J. Li, G. Colak, Y. Chen, C. Xia, C. Peng, H. Ruan, M. Kirkey, D. Wang, L.M. Jensen, O.K. Kwon, S. Lee, S.D. Pletcher, M. Tan, D.B. Lombard, K.P. White, H. Zhao, J. Li, R.G. Roeder, X. Yang, Y. Zhao, Metabolic Regulation of Gene Expression by Histone Lysine beta-Hydroxybutyrylation. Mol cell 62, 194–206 (2016).

6. L.T. Izzo, K.E. Wellen, Histone lactylation links metabolism and gene regulation. Nature 574, 492–493 (2019).

7. L.A. Farrelly, R.E. Thompson, S. Zhao, A.E. Lepack, Y. Lyu, N.V. Bhanu, B. Zhang, Y.E. Loh, A. Ramakrishnan, K.C. Vadodaria, K.J. Heard, G. Erikson, T. Nakadai, R.M. Bastle, B.J. Lukasak, H. Zebroski, 3rd, N. Alenina, M. Bader, O. Berton, R.G. Roeder, H. Molina, F.H. Gage, L. Shen, B.A. Garcia, H. Li, T.W. Muir, I. Maze, Histone serotonylation is a permissive modification that enhances TFIID binding to H3K4me3. Nature 567, 535–539 (2019).

8. Y. Li, B.R. Sabari, T. Panchenko, H. Wen, D. Zhao, H. Guan, L. Wan, H. Huang, Z. Tang, Y. Zhao, R.G. Roeder, X. Shi, C.D. Allis, H. Li, Molecular Coupling of Histone Crotonylation and Active Transcription by AF9 YEATS Domain. Mol cell 62, 181–193 (2016).

9. D.J. Walther, M. Bader, Aunique central tryptophan hydroxylase isoform. Biochem Pharmacol 66, 1673–1680 (2003).

10. A. Dahlstrom, K. Fuxe, Localization of monoamines in the lower brain stem. Experientia 20, 398–399 (1964).

11. R. Hummerich, P. Schloss, Serotonin--more than a neurotransmitter: transglutaminase-mediated serotonylation of C6 glioma cells and fibronectin. Neurochem Int 57, 67–75 (2010).

12. D.J. Walther, J.U. Peter, S. Winter, M. Holtje, N. Paulmann, M. Grohmann, J. Vowinckel, V. Alamo-Bethencourt, C.S. Wilhelm, G. Ahnert-Hilger, M. Bader, Serotonylation of small GTPases is a signal transduction pathway that triggers platelet alpha-granule release. Cell 115, 851–862 (2003).

13. Y. Dai, N.L. Dudek, T.B. Patel, N.A. Muma, Transglutaminase-catalyzed transamidation: a novel mechanism for Rac1 activation by 5-hydroxytryptamine2A receptor stimulation. J Pharmacol Exp Ther 326, 153–162 (2008).

14. E. Ballestar, C. Abad, L. Franco, Core histones are glutaminyl substrates for tissue transglutaminase. J Biol Chem 271, 18817–18824 (1996).

15. J. Wysocka, T. Swigut, T.A. Milne, Y. Dou, X. Zhang, A.L. Burlingame, R.G. Roeder, A.H. Brivanlou, C.D. Allis, WDR5 associates with histone H3 methylated at K4 and is essential for H3 K4 methylation and vertebrate development. Cell 121, 859–872 (2005).

16. Y.S. Ang, S.Y. Tsai, D.F. Lee, J. Monk, J. Su, K. Ratnakumar, J. Ding, Y. Ge, H. Darr, B. Chang, J. Wang, M. Rendl, E. Bernstein, C. Schaniel, I.R. Lemischka, Wdr5 mediates self-renewal and reprogramming via the embryonic stem cell core transcriptional network. Cell 145, 183–197 (2011).

17. C. Lin, Y. Wang, Y. Wang, S. Zhang, L. Yu, C. Guo, H. Xu, Transcriptional and posttranscriptional regulation of HOXA13 by lncRNA HOTTIP facilitates tumorigenesis and metastasis in esophageal squamous carcinoma cells. Oncogene 36, 5392–5406 (2017).

18. T. Nakagawa, Y. Xiong, X-linked mental retardation gene CUL4B targets ubiquitylation of H3K4 methyltransferase component WDR5 and regulates neuronal gene expression. Mol cell 43, 381–391 (2011).

19. J.F. Couture, E. Collazo, R.C. Trievel, Molecular recognition of histone H3 by the WD40 protein WDR5. Nat Struct Mol Biol 13, 698–703 (2006).

20. A. Schuetz, A. Allali-Hassani, F. Martin, P. Loppnau, M. Vedadi, A. Bochkarev, A.N. Plotnikov, C.H. Arrowsmith, J. Min, Structural basis for molecular recognition and presentation of histone H3 by WDR5. EMBO J 25, 4245–4252 (2006).

21. Z. Han, L. Guo, H. Wang, Y. Shen, X.W. Deng, J. Chai, Structural basis for the specific recognition of methylated histone H3 lysine 4 by the WD-40 protein WDR5. Mol cell 22, 137–144 (2006).

22. A.J. Ruthenburg, W. Wang, D.M. Graybosch, H. Li, C.D. Allis, D.J. Patel, G.L. Verdine, Histone H3 recognition and presentation by the WDR5 module of the MLL1 complex. Nat Struct Mol Biol 13, 704–712 (2006).

23. Y. Sun, J.L. Bell, D. Carter, S. Gherardi, R.C. Poulos, G. Milazzo, J.W. Wong, R. Al-Awar, A.E. Tee, P.Y. Liu, B. Liu, B. Atmadibrata, M. Wong, T. Trahair, Q. Zhao,J. M. Shohet, Y. Haupt, J.H. Schulte, P.J. Brown, C.H. Arrowsmith, M. Vedadi, L. MacKenzie, S. Huttelmaier, G. Perini, G.M. Marshall, A. Braithwaite, T. Liu, WDR5 Supports an N-Myc Transcriptional Complex That Drives a Protumorigenic Gene Expression Signature in Neuroblastoma. Cancer Res 75, 5143–5154 (2015).

24. L.R. Thomas, C.M. Adams, J. Wang, A.M. Weissmiller, J. Creighton, S.L. Lorey, Q. Liu, S.W. Fesik, C.M. Eischen, W.P. Tansey, Interaction of the oncoprotein transcription factor MYC with its chromatin cofactor WDR5 is essential for tumor maintenance. Proc Natl Acad Sci USA 116, 25260–25268 (2019).

25. R. Huang, N.K. Cheung, J. Vider, I.Y. Cheung, W.L. Gerald, S.K. Tickoo, E.C. Holland, R.G. Blasberg, MYCN and MYC regulate tumor proliferation and tumorigenesis directly through BMI1 in human neuroblastomas. FASEB J 25, 4138–4149 (2011).

26. D. Zhang, F. Wang, Y. Pang, E. Zhao, S. Zhu, F. Chen, H. Cui, ALG2 regulates glioblastoma cell proliferation, migration and tumorigenicity. Biochem Biophys Res Commun 486, 300–306 (2017).

27. J.R. Lee, J.L. Roh, S.M. Lee, Y. Park, K.J. Cho, S.H. Choi, S.Y. Nam, S.Y. Kim, Overexpression of glutathione peroxidase 1 predicts poor prognosis in oral squamous cell carcinoma. J Cancer Res Clin Oncol 143, 2257–2265 (2017).

28. B.K. Neilsen, B. Chakraborty, J.L. McCall, D.E. Frodyma, R.L. Sleightholm, K.W. Fisher, R.E. Lewis, WDR5 supports colon cancer cells by promoting methylation of H3K4 and suppressing DNA damage. BMC Cancer 18, 673 (2018).

29. R. Malek, R.P. Gajula, R.D. Williams, B. Nghiem, B.W. Simons, K. Nugent, H. Wang, K. Taparra, G. Lemtiri-Chlieh, A.R. Yoon, L. True, S.S. An, T.L. DeWeese, A.E. Ross, E.M. Schaeffer, K.J. Pienta, P.J. Hurley, C. Morrissey, P.T. Tran, TWIST1-WDR5-Hottip Regulates Hoxa9 Chromatin to Facilitate Prostate Cancer Metastasis. Cancer Res 77, 3181–3193 (2017).

30. W. Sun, F. Guo, M. Liu, Up-regulated WDR5 promotes gastric cancer formation by induced cyclin D1 expression. J Cell Biochem 119, 3304–3316 (2018).

31. S. Punzi, C. Balestrieri, C. D’Alesio, D. Bossi, G.I. Dellino, E. Gatti, G. Pruneri, C. Criscitiello, G. Lovati, M. Meliksetyan, A. Carugo, G. Curigliano, G. Natoli, P.G. Pelicci, L. Lanfrancone, WDR5 inhibition halts metastasis dissemination by repressing the mesenchymal phenotype of breast cancer cells. Breast Cancer Res 21, 123 (2019).

32. H. Chen, B. Lorton, V. Gupta, D. Shechter, A TGFbeta-PRMT5-MEP50 axis regulates cancer cell invasion through histone H3 and H4 arginine methylation coupled transcriptional activation and repression. Oncogene 36, 373–386 (2017).

33. X. Chen, W. Xie, P. Gu, Q. Cai, B. Wang, Y. Xie, W. Dong, W. He, G. Zhong, T. Lin, J. Huang, Upregulated WDR5 promotes proliferation, self-renewal and chemoresistance in bladder cancer via mediating H3K4 trimethylation. Sci Rep 5, 8293 (2015).

34. S.M. Lauberth, T. Nakayama, X. Wu, A.L. Ferris, Z. Tang, S.H. Hughes, R.G. Roeder, H3K4me3 interactions with TAF3 regulate preinitiation complex assembly and selective gene activation. Cell 152, 1021–1036 (2013).

35. J. Wysocka, T. Swigut, H. Xiao, T.A. Milne, S.Y. Kwon, J. Landry, M. Kauer, A.J. Tackett, B.T. Chait, P. Badenhorst, C. Wu, C.D. Allis, A PHD finger of NURF couples histone H3 lysine 4 trimethylation with chromatin remodelling. Nature 442, 86–90 (2006).

36. W. Wang, Z. Chen, Z. Mao, H. Zhang, X. Ding, S. Chen, X. Zhang, R. Xu, B. Zhu, Nucleolar protein Spindlin1 recognizes H3K4 methylation and stimulates the expression of rRNA genes. EMBO Rep 12, 1160–1166 (2011).

37. X. Su, G. Zhu, X. Ding, S.Y. Lee, Y. Dou, B. Zhu, W. Wu, H. Li, Molecular basis underlying histone H3 lysine-arginine methylation pattern readout by Spin/Ssty repeats of Spindlin1. Genes Dev 28, 622–636 (2014).

38. Z. Otwinowski, W. Minor, Processing of X-Ray Diffraction Data Collected in Oscillation Mode. Methods Enzymol 276, 307–326 (1997).

39. A.J. McCoy, Solving structures of protein complexes by molecular replacement with Phaser. Acta Crystallogr D Biol Crystallogr 63, 32–41 (2007).

40. P.D. Adams, R. W. Grosse-Kunstleve, L.W. Hung, T.R. Ioerger, A.J. McCoy, N.W. Moriarty, R.J. Read, J.C. Sacchettini, N.K. Sauter, T.C. Terwilliger, PHENIX: building new software for automated crystallographic structure determination. Acta Crystallogr D Biol Crystallogr 58, 1948–1954 (2002).

41. A.A. Vagin, R.A. Steiner, A.A. Lebedev, L. Potterton, S. McNicholas, F. Long, G.N. Murshudov, REFMAC5 dictionary: organization of prior chemical knowledge and guidelines for its use. Acta Crystallogr D Biol Crystallogr 60, 2184–2195 (2004).

42. P. Emsley, K. Cowtan, Coot: model-building tools for molecular graphics. Acta Crystallogr D Biol Crystallogr 60, 2126–2132 (2004).

