## supplemental tables and figures for "Structural insights into the recognition of histone H3Q5 serotonylation by WDR5"

**Supplementary information**

**Supplementary Figures**

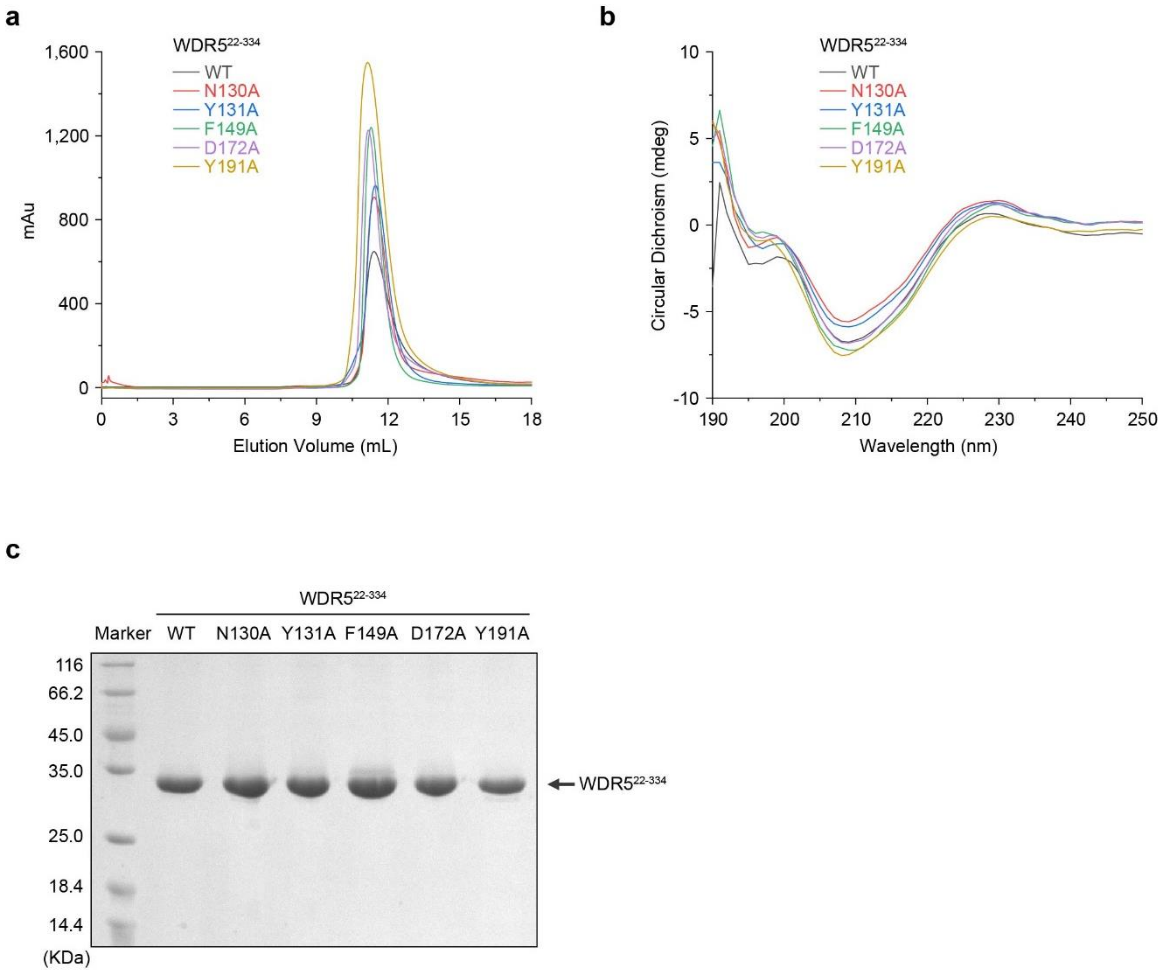


**Supplementary Fig. 1**

**a-b,** Size-Exclusion chromatography (**a**) and CD spectrum (**b**) results of WT WDR5$^{22-334}$ and five putative H3Q5ser-binding-deficient WDR5 mutant proteins, *i.e.*, N130A, Y131A, F149A, D172A, and Y191A.

**c**, Representative of purified WT and aforementioned mutant WDR5$^{22-334}$ proteins detected by SDS-PAGE.

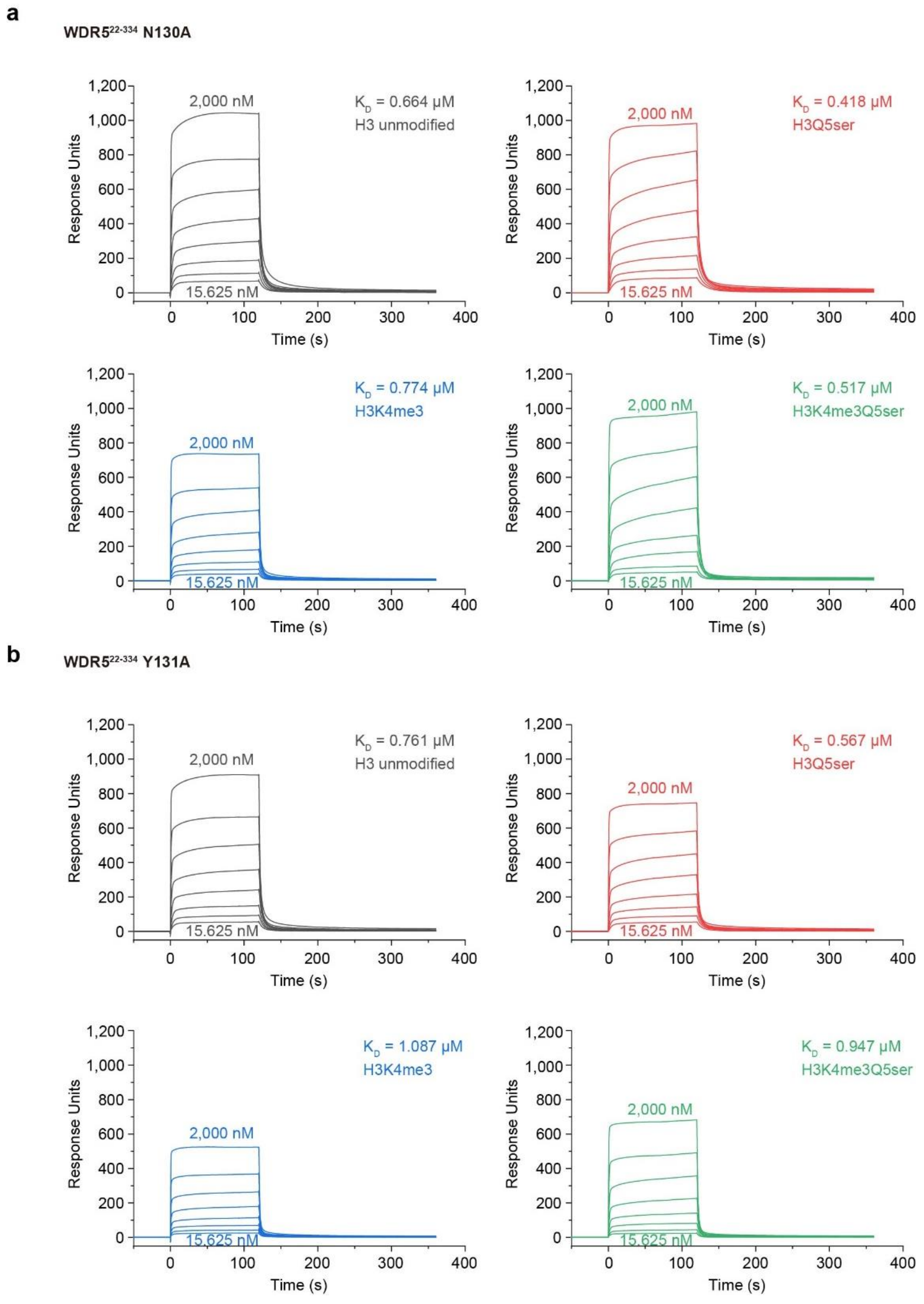


**Supplementary Fig. 2**

**a-b**, Dissociation constants of WDR5$^{22\text{-}334}$ N130A (**a**) and Y131A (**b**) with different H3 peptides determined by SPR assay.

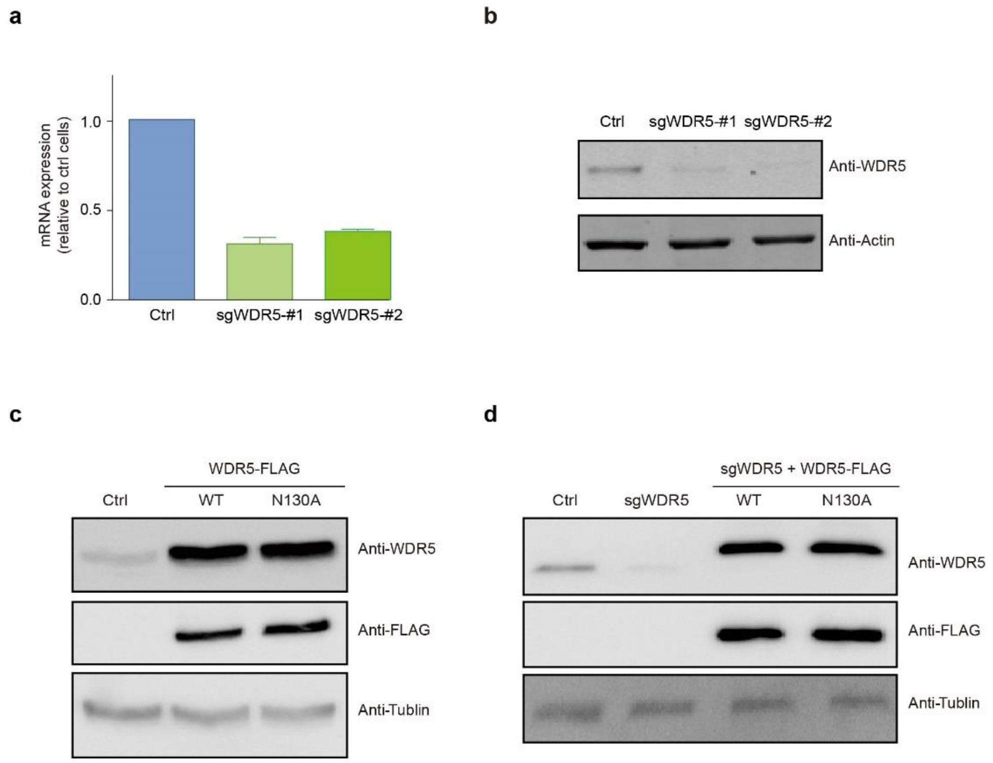


**Supplementary Fig. 3**

**a-b**, mRNA (**a**) and protein (**b**) levels of WDR5 in CRIPSR-Cas9-mediated WDR5 knockout (WDR5$^{-/-}$) SK-N-SH cells determined by qPCR and Western blotting, respectively.

**c-d**, Overexpression of wild type or N130A mutant WDR5 in control (**c**) or WDR5$^{-/-}$ (**d**) SK-N-SH cells.

**Supplementary Tables**

**Supplementary Table S1. Sequences of various synthesized H3 peptides**

| | |
|---|---|
| H3 unmodified-biotin | ARTKQTARKSTGGK-biotin |
| H3K4me3-biotin | ARTK (me3) QTARKSTGGK-biotin |
| H3Q5ser-biotin | ARTKQ (ser) TARKSTGGK-biotin |
| H3K4me3Q5ser-biotin | ARTK (me3) Q (ser) TARKSTGGK-biotin |
| H3Q5ser | ARTKQ (ser) TARKSTGGK |
| H3K4me3Q5ser | ARTK (me3) Q (ser) TARKSTGGK |

**Supplementary Table 2. Sequence of the primers used for qPCR**

| | |
|---|---|
| PDCD6-F | TCCAGAGGGTCGATAAAGACA |
| PDCD6-R | TTCTGCCAGTCCGTGATGT |
| GPX1-F | GCGGGGCAAGGTACTACTTA |
| GPX1-R | CTCTTCGTTCTTGGCGTTCT |
| C-MYC-F | GGCTCCTGGCAAAAGGTCA |
| C-MYC-R | AGTTGTGCTGATGTGTGGAGA |
| WDR5-F | AATTCAGCCCGAATGGAGAGT |
| WDR5-R | AGGCTACATCGGATATTCCCAG |
| Actin-F | CATGTACGTTGCTATCCAGGC |
| Actin-R | CTCCTTAATGTCACGCACGAT |

**Supplementary Table 3. Sequence of the primers used in ChIP-qPCR assay**

| | |
|---|---|
| PDCD6-F | GGAAAGACCCAGGGATCACA |
| PDCD6-R | CCTTCGTGCTGGACACAGTT |
| GPX1-F | TGGAGAGGCCATGACTCCAA |
| GPX1-R | AGGCAATGGTTGACTGCCTA |
| C-MYC-F | TCCACAAGCTCTCCACTTGC |
| C-MYC-R | GCCGGGTTTGGGAGAAATCA |
